# Overestimation of the prevalence of eosinophilic colitis with reliance on a single billing code

**DOI:** 10.1101/414557

**Authors:** AB Muir, ET Jensen, JB Wechsler, P Menard-Katcher, GW Falk, SS Aceves, GT Furuta, ES Dellon, ME Rothenberg, JM Spergel

## Abstract

Eosinophilic colitis (EC) is a rare disorder characterized by eosinophilic inflammation of the colon causing diarrhea, bloody stool, nausea, constipation and abdominal pain. The Consortium for Eosinophilic Gastrointestinal Disease Research sought to undertake the first multi-center study of the EC population, but faced challenges with meeting enrollment goals that were based on initial estimates. To understand the reason for this, we performed chart review of patients with ICD codes for EC at 8 tertiary care centers. Chart review revealed that the isolated use of ICD codes overestimated EC rates in the pediatric population.

## Introduction

Eosinophilic colitis (EC) is a chronic disease characterized by eosinophil-predominant inflammation of the colon. Studies probing claims data suggest that the prevalence of EC is 1.5-2.4/100,000^1,2^. International Classification of Disease (ICD) and Systematized Nomenclature of Medicine (SNOMED) codes are classification systems that can be used in the studies of administrated claims data. The Consortium for Eosinophilic Gastrointestinal Disease Research (CEGIR), part of NIH’s Rare Diseases Clinical Research Network^3^, planned a prospective multi-center longitudinal study with a projected enrollment of 150 patients over the course of 4 years. However, only 31 patients have been enrolled over the course of the 3-year enrollment period to date.

In order to diagnose EC, other inflammatory (inflammatory bowel disease (IBD), allergic proctocolitis of infancy, food protein induced enterocolitis (FPIES)) or infectious (parasite) etiologies must be ruled out. Differentiating between these conditions using claims data alone can be challenging as EC and allergic proctocolitis of infancy share the same ICD 9/10 codes and EC and FPIES share ICD9 codes. We therefore hypothesized that ambiguity in ICD coding has led to overestimation of the rates of EC.

## Methods

We performed a retrospective study to assess the prevalence of EC at 8 institutions (5 pediatric and 3 adult hospitals) over the last 3-10 years (Table 1). Electronic medical records were searched for all ICD codes for EC (ICD9:558.42; ICD10:K52.82). It is important to note that these codes can be used for other diseases that cause eosinophilia in the colon. Charts were evaluated for co-morbid conditions that could provide alternative explanations for colonic eosinophilia (inflammatory bowel disease, food-protein induced enterocolitis, allergic proctocolitis of infancy, malignancy, and parasitic infection). Pathology reports were reviewed at each site to confirm eosinophilic infiltrate of the colon.

**Table 1:** Confirmed cases of EC at each institution

| **Hospital Type** | **Institution** | **Years of Chart Review** | **ICD9/10 coded EC patients (n)** | **EC Diagnosis confirmed (n)** |
| --- | --- | --- | --- | --- |
| Pediatric | The Children's Hospital of Philadelphia | 4 | 44 | 1 |
| Pediatric | Cincinnati Children's Hospital | 10 | 228 | 29 |
| Pediatric | Rady Children's Hospital San Diego | 3 | 32 | 2 |
| Pediatric | Children's Hospital of Colorado | 6 | 185 | 5 |
| Pediatric | Lurie Children's Hospital of Chicago | 4 | 12 | 1 |
|  | **Total Pediatric** |  | **501** | **38** |
| Adult | Hospital of the University of Pennsylvania | 8 | 9 | 3 |
| Adult | University of North Carolina Hospital | 3 | 10 | 7 |
| Adult | University of Colorado Hospital | 8 | 1 | 1 |
|  | **Total Adult** |  | **20** | **11** |

## Results

In 5 pediatric hospitals, 510 patients had an ICD9/10 code for EC, but just 38 out of 501 (7%) of the cases met clinical and histopathologic features suggestive of EC. In 3 adult hospitals, there were only 20 patients with an ICD code for EC; 11 out of 20 (55%) had confirmed EC after chart review (Table 1). This stark difference between the pediatric and adult centers was in large part due to overlapping ICD codes for more common pediatric diseases, specifically food-protein induced enterocolitis (FPIES) and allergic colitis of infancy both of which shared ICD-9/10 codes with eosinophilic colitis.

## Discussion

These data underscore the importance of establishing both clinical-histopathologic definitions and specific ICD codes for EC. Collins et al. proposed that in addition to exclusion of other inflammatory conditions, 100 eos/HPF in cecum/ascending colon, 84/HPF in transverse/descending colon, or 64/HPF in rectosigmoid^4^ represent pathologic eosinophilia of the colon. However, these definitions have yet to be clinically validated or adopted by the pathology or gastroenterology communities, and are currently used only for research purposes; indeed arriving at a consensus criteria is a goal of CEGIR. The lack of diagnostic criteria presents clear limitations for the utilization of the ICD-9-10 code in identifying true cases of EC.

While use of claims databases has distinct advantages such as the ability to capture a large population in geographically diverse areas, there is the risk of misclassification due to lack of granularity of patient data. For example, in the studies assessing EC prevalence, pathology reports were not available in the administrative/insurance claim databases^1,2^. These studies defined an EC case according to the presence of one more instances of ICD-9/10 codes for EC, and with exclusion of cases with certain co-morbid conditions that could explain eosinophilia (e.g. inflammatory bowel disease)^1,2^. The positive predictive value of using administrative claims data for identifying EC cases may be further improved by following the approach of case ascertainment in other GI conditions. For example, for inflammatory bowel disease (IBD), ICD codes alone were not sufficient to accurately describe the population, but when combined with IBD-specific pharmaceutical claims, accuracy was improved^5^. Likewise, in Barrett’s esophagus, ICD codes overestimate prevalence with only 27% of coded patients having disease^6^. In both IBD and Barrett’s there are diagnostic criteria and treatment algorithms that complement ICD data. However, in the case of EC there is broad symptomatology, no therapeutic or diagnostic guidelines, and non-specific ICD codes. Ultimately, administrative/insurance claims databases for purposes of examining eosinophilic gastrointestinal disorders will necessitate not only identifying ICD codes, but also procedure codes (for colonoscopy and biopsy), pathology interpretation codes, exclusion of codes for competing conditions, and perhaps exclusion of infants under the age of 1 to exclude proctocolitis of infancy.

In summary, we have found that EC is less common than previously described suggesting 10x fold over-diagnosis of EC using pediatric coding. In adults, we also found a paucity of cases, although overdiagnosis using ICD coding is to a lesser extent. Establishing clinical-histopathologic guidelines and distinct ICD codes for EC will allow for better estimation of disease prevalence and will inform future clinical and translational studies involving EGIDs.

